# In vitro and in vivo Antioxidant potential of aqueous and hydroethanolic extracts leaves of *Dracaena deisteliana* Eng. (*Dracacenaceae*)

**DOI:** 10.1101/2023.01.24.525426

**Authors:** Huguette Bocanestine Laure Feudjio, Marie Stéphanie Chekem Goka, Josias Djeguemtar, Gabriel Tchuente Kamsu, Jean Baptiste Sokoudjou, Louis-Claire Ndel Famen, Norbert Kodjio, Donatien Gatsing

## Abstract

Enterobacteria such as those of the genus *Salmonella* express the oxyR gene that codes for several proteins that allow it to resist free radicals. This resistance of *Salmonella* is often at the origin of an overproduction of these free radicals that can lead to oxidative stress. To investigate the antioxidant activity in vitro (aqueous and hydroethanolic extracts in vivo (of the 55% hydroethanolic extract) of Dracaena deisteliana leaves in albino rats of Wistar strain previously infected with the Salmonella strain Typhi ATCC 6539. The *in vitro* antioxidant activity of these extracts was determined by studying their anti-radical power with DPPH•, then their iron-reducing power and determining their flavonoids and total phenols content. *In vivo* antioxidant activity was determined in plasma and tissues of albino rats of wistar strain aged between 8 and 10 weeks previously infected with *Salmonella* typhi ATCC 6539. These infected rats concurrently received daily doses of *Dracaena deisteliana* extract (10, 20 and 51.84 mg/kg) or ciprofloxacin (14 mg/kg) as positive control, for 15 days. At the end of the treatment period, the animals were sacrificed and blood, liver, kidney, heart, lung, spleen testis and ovaris were collected for evaluation of antioxidants parameters, which included malondialdehyde, nitric oxide, catalase and peroxidase, as well as biological responses. Regarding *in vitro* antioxidant tests, the 65% hydroethanolic extract showed an anti-radical activity with DPPH• superior to that of all hydroethanolic extracts at 100 μg/ml. However, the infused showed lower antiradical activity than all extracts at 12.5 and 200 μg/ml concentrations. The 55% hydroethanolic extract had the best IC_50_ of (11.99 μg/ml). The iron reducing power of this extract was higher than the other extracts at 200 μg/ml. The highest levels of flavonoids and total phenols were obtained respectively with the 55% and 95% hydroethanolic extract. The hydroethanol extract of *Dracaena deisteliana* (10, 20 and 51.84 mg/kg) cured the infected rats between the 9th and 13th day of treatment. The extract also significantly reduced (p < 0.05) blood malondialdehyde and nitric oxide levels, and significantly increased (p >0.05) the activity of catalase and peroxidase in the infected rats. The results suggest that leaves extract of *Dracaena deisteliana* contains antisalmonella and antioxidant substances, which could be used for the treatment of typhoid fever and another salmonellosis. In addition, 55% hydroethanolic extract of this plant possesses antisalmonella activity and reduces the state of oxidative stress caused by S. typhi during rat’s infection.

## INTRODUCTION

During microbial infections, macrophages produce free radicals in order to destroy microorganisms within phagosomes. However, some enterobacteriaceae such as those of the genus *Salmonella* express the oxyR gene which codes for several proteins allowing it to resist these free radicals [1, 2]. This resistance of *Salmonella* is often the cause of an overproduction of free radicals that can lead to oxidative stress in Salmonella infected patients [3,4]. In addition, during a *salmonella* infection, or following the exposure of the body to exogenous toxins, the production of free radicals such as superoxide anion and nitric oxide (O^2−^, NO.), athough controlled by antioxidant defense systems under normal physiological conditions, what can increase and generate oxidative stress[5]. This state of oxidative stress is the direct cause of various pathological conditions such as aging and cancer: it is the indirect cause of lipid peroxidation in food. In all cases, the risk is increased with the accumulation of these molecules in the body resulting in a radical chain reaction that degrades vital biological molecules, namely DNA, lipids, proteins and carbohydrates [6, 7, 8]. Frequent exposure to high concentrations of free radicals can lead to direct damage to biological molecules (protein oxidation and nitration, lipid peroxidation, DNA oxidation), but also secondary lesions due to the cytotoxic and mutagenic nature of the metabolites released during lipid oxidation [9,10].

Thus, treatment sources to combat both *Salmonella* infection and oxidative stress would improve the management of patients suffering from these infections [11, 12, 13, 14, 15]. Since ancient times, medicinal plants have always been part of the basic knowledge of all human societies [16]. In this era of rapid advances in medical technology, herbal preparations used in “alternative or complementary medicine” are gaining a lot of popularity [17], and increased interest in their use has encouraged more detailed studies of plant resources [18]. Previous studies have found that *Dracaena deisteliana* leaves commonly used in combination with *Senecio biafrae* leaves and stems help fight stomach upset and infertility in women [19]. The roots of this plant are used to treat toothache problems [20]. No studies on *in vitro* and *in vivo* antioxidant activities have yet been evaluated, hence the importance of this work.

The purpose of this work was to make a study to provide information on the *in vitro* antioxidant activities of the aqueous and hydroethanolic extracts of the leaves of *Dracaena deisteliana* and to evaluate the *in vivo* antioxidant potential of the 55% hydroethanolic extract of *Dracaena deisteliana* leaves for oxidative stress induced by *Salmonella* Typhi (ATCC 6539) in albino rats of wistar strain.

## MATERIALS AND METHODS

### Plant material

The fresh leaves sample of *Dracaena deisteliana* was harvested in January 2020 at Campus of Dschang University (Department of Menoua, Western Region of Cameroon). The identification of this plant was made at the National Herbarium of Cameroon (Yaoundé) compared to the sample registered under the number 53011HNC.

### Bacteria strain

Stock cultures of S. Typhi (ATCC 6539) used in this study were obtained from Centre Pasteur of Cameroon. Their pure cultures were maintained in Muller-Hinton agar and stored at 4°C.

### Experimental animals

Young, healthy Wistar rats aged 8 to 10 weeks of each sex were bred at the University of Dschang animal house. Animal housing and *in vivo* experiments were carried out following the guidelines of the European Union on Animal Care (CEE Council 86/609)15 that were adopted by the Institutional Committee of the Ministry of Scientific Research and Innovation of Cameroon. They were housed under a natural temperature and 12 hours dark/light cycle. The animals were fed with standard diet and received water *ad libitum*. All animal procedures were performed after approval by the University of Dschang-Cameroon Ethics Committee (Project N° BCH1202/FS/UDs/2018).

### Preparation of extracts

The fresh leaves of *Dracaena deisteliana* was air dried for three weeks at room temperature (24 to 27°C) until constant weight; then mashed. The obtained powder was used for the preparation of hydroethanolic extracts (95%, 85%, 75%, 65%, 55% and 45%) and aqueous extracts (infusion, decoction, maceration). Aqueous extracts were prepared according to the methods described by [15]. Whith some modifications, while hydroethanolic extract were obtained by macerating 50 g of powder in 500 ml in hydroethanolic at defferent concentrations (95%, 85%, 75%, 65%, 55% and 45%). After 48 hours, these macerates were filtered using Whatman N°1 paper. Each hydroethanolic extract was evaporated at 40 °C using rotary evaporator (BUCHI R-200). The filtrates were dried at 45 °C in a ventilated oven (Memmert) for seven days in order to completely evaporate the rest of solvent.

### *In vitro* antioxidant activities of aqueous and hydroethanolique extracts of *Dracaena deisteliana* leaves

#### 2, 2-Diphenyl-picryl-hydrazyl (DPPH) radical scavenging assay

The free radical scavenging activities of *Dracaena deisteliana* leaves extracts were evaluated using the DPPH assay method as described by [21] and [22]. Briefly, the extract (2000 μg/ml) was twofold serially diluted with methanoln, hence 100 μl of diluted extract were mixed with 900 μl of 0.3 mM 2,2-diphenyl-1-picrylhydrazyl (DPPH) methanol solution to give a final extract concentration of 12.5, 25, 50, 100 and 200 μg/ml. After 30 min of incubation in the dark at room temperature, the optical densities were measured at 517 nm. Ascorbic acid (Vitamin C) was used as control. Each assay was done in triplicate and the results, recorded as mean ± standard deviation of three findings, were illustrated in tabular form. The percentage of DPPH scavenging activity (%DPPH) was calculated according to the following equation:

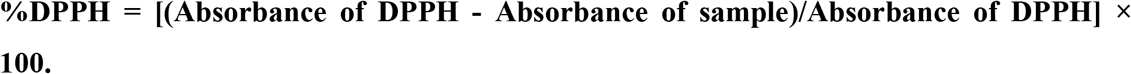

The IC_50_ is automatically determined from the percentages of inhibitions obtained from the absorbances of DPPH• according to the different concentrations of extracts and vitamin C.

#### Ferric reducing antioxidant power (FRAP) assay

The ferric reduction potential (conversion potential of Fe3+ to Fe2+) of extracts was determined according to the method described by [23]. To do this, 1 ml of extract at different concentrations (200, 100, 50, 25 and 12.5 μg/ml) was mixed with 2.5 ml of a 0.2 M phosphate buffer solution (pH 6.6) and 2.5 ml of a solution of potassium ferricyanide K_3_Fe(CN)_6_ at 1%. The mixture was incubated in a water bath at 50°C for 20 min; then 2.5 ml of 10% trichloroacetic acid was added to stop the reaction, and tubes were centrifuged at 3000 rpm for 10 min. An aliquot (2.5 ml) of supernatant was combined with 2.5 ml of distilled water and 0.5 ml of 0.1% methanolic FeCl3 solution. The absorbance of the reaction medium was read at 700 nm against a similarly prepared control, replacing the extract with distilled water. The positive control was represented by a solution of a standard antioxidant, ascorbic acid whose absorbance was measured under the same conditions as the samples. Increased absorbance of the reaction mixture indicates higher reduction capacity of the tested extracts [24, 25].

#### Evaluation of total phenolic contents

The total phenol content was determined by the method described by Ramde- [26]. The Folin-Ciocateu method was used for the quantitative determination of total phenolic compounds. The reaction mixture consisted of 0.02 ml of extract (2000 μg/ml), 1.38 ml of distilled water, 0.02 ml of 2N Folin-ciocalteu reagent and 0.4 ml of a 20% sodium carbonate solution. The mixture was then incubated at 40 °C for 20 min and the absorbance was measured at 760 nm. The blank was prepared with distilled water instead of plant extracts. Each test was performed in triplicate and the results were expressed as milligrams of Equivalents Gallic Acid (mgEGA) per gram of extract using a standard calibration curve of gallic acid (0.2 g/l; 0-70 μl).

#### Evaluation of total flavonoids contents

The flavonoid content of the extracts was determined using the aluminum trichloride colorimetric method [27]. The determination of total flavonoids content was done using the colorimetric aluminum chloride method. In fact, 100 μl of extract (2 mg/ml) were mixed with 1.49 ml of distilled water and 30 μl of NaNO2 (5%). After 5 min incubation at room temperature, 30 μl of AlCl3 (10%) was added and the mixture was reincubated for 6 min; then 200 μl of sodium hydroxide (0.1 M) and 240 μl of distilled water were added. The solution was well mixed and the absorbance was read at 510 nm using a spectrophotometer. Each test was performed in triplicate and the results were expressed as milligrams of Equivalents Catechin (mgECat) per gram of extract using a standard calibration curve of Catechin (0.1 g/l; 0-70 μl).

### *In vivo* antioxidant activity of the 55% hydroethanolic extract of the leaves of *Dracaena deisteliana*

The animals were infected with *Salmonella* Typhi and then treated; during treatment, we studied the evolution of the bacterial load in the animals’ blood. For this study, which lasted 15 days, 48 albino rats (male and female) of Wistar strain aged 8 to 10 weeks were used. The animals were selected by sorting after culture of their blood on *Salmonella*-Shigella agar.

#### Typhoid fever induction and treatment

Antityphic activity was determined according to the protocol described by [14], with some modifications was used. Forty-eight male and female Albino Wistar rats aged 08 to 10 weeks were divided into 12 groups of 4 animals of each sex, including 6 male groups and 6 female groups. The selected animals were acclimatized for a week. With the exception of animals in group 1 of each sex (uninfected and untreated), all animals in the other groups (2-6) were infected. They received a single dose (1 mL) of a suspension of 1.5×10^8^ CFU of S. typhi (ATCC 6539) orally. Infection monitoring in animals was performed by blood culture with colony counting on Salmonella-Shigella agar and converted to salmonella CFU per milliliter of blood. The success of the infection was demonstrated when the concentration of bacteria in the blood was greater than 4×10^5^ CFU / ml of blood, then by the excretion of watery stools, the presence of mucus in the stool, the reduction of activity and the exponential increase in the systemic load of S. typhi in rats [24]. Each animal in each group was housed in its own cage and these animals were treated as follows: group 1 (neutral control group) received distilled water; group 2 (typhoid control group, infected and untreated) received distilled water; group 3 (positive control group) received ciprofloxacin (14 mg/kg); groups 4, 5 and 6 (test groups) received a hydroethanolic extract 55% of *Dracaena deisteliana* (10, 20 and 51.84 mg/Kg respectively). Dose 51.84 mg/kg body weight were obtained from the daily dose of the traditional practitioner, the dose of 10 mg/kg body weight was obtained from the MIC of the extract 20 mg/kg body weight is double the dose of the obtained from the MIC. Then, every 2 days during the experimental period, blood was taken from the caudal vein located on the tail of the rats and introduced into heparinated tubes. Then diluted to 1/10th with physiological water, 50 μl of the mixture was taken and introduced into the sterile petri dishes previously filled with Salmonella-Shigella agar for the enumeration of the bacterial load. The decrease in the bacterial load in the blood indicated the effectiveness of the treatment. The test was completed when no more than two colonies of salmonella were found in the animals’ blood after the blood culture. Food consumption as well as body weight of animals were recorded daily as described by [12].

#### Animal sacrifice

The day before the end of treatment, the animals were subjected to a 12 hour fast at the end of which the urine was collected in the previously washed cages and lined with wire mesh. Subsequently, the animals were anesthetized by intraperitoneal injection of Diazepam and Ketamine (0.2+0.1) mL; then placed in supine position on a board, and all four limbs were immobilized to allow easy access to the abdomen. A dissection was performed on the abdomen and the blood was collected by cardiac puncture and then first introduced into two tubes (for each animal) each containing an anticoagulant (EDTA); one for the determination of hematological parameters and the other for the preparation of plasma. This plasma was obtained by centrifugation at 3000 rpm for 15 minutes after letting stand for 4 hours of time. Organs such as the heart, kidneys, liver, lung and spleen were removed and the homogenates of the heart, kidneys, liver, lungs and spleen were prepared in a physiological water solution, at the rate of 15 g of tissue per 100 ml of buffer, then centrifuged at 3000 rpm for 15 min. The resulting crush was also centrifuged for 15 minutes at 3000 rpm and the supernatant was recovered for the different dosages. The plasmas and homogenates thus obtained were kept at −18 °C for the determination of antioxidant parameters.

#### Determination of nitric oxide (NO)

Nitric oxide content in plasma and tissue homogenates was estimated from the accumulation of nitrite (NO_2_^−^) using Griess’ reagent, as described by [28]. With some modifications. Absorption of the chromophore during ionization of nitrite with sulfanilamide coupled with naphthylethylenediamine (NED) was read at 520 nm. Three hundred and forty microliters of 1% sulfanilamide (prepared in 5% orthophosphoric acid) was introduced into 340 μL of plasma and homogenates. The resulting mixture was homogenized and left in the dark for 5 minutes at room temperature. Next, 340 μL of 0.1% NED was added to the reaction medium and the whole was left once more in the dark for 5 minutes. Optical densities were read against the control at the wavelength of 520 nm and the results were expressed in terms of micromoles of NO per gram of tissue or per milliliter of blood as a function of the standard equation of NO (y = 0.0563x + 0.1077).

#### Determination of malondialdehyde (MDA)

The lipid peroxidation index by the measurement of malondialdehyde (MDA) was measured in tissues using thiobarbituric acid (TBA) according to the method of [29], with certain modifications, malondialdehyde is one of the final products of the decomposition of polyunsaturated fatty acids (PUFAs) under the effect of free radicals released during stress. In a hot acidic medium (pH 2 to 3; 100°C), an MDA molecule condenses with two thiobarbituric molecules (TBA) to form a pink colored complex (reading at 532 nm). Five hundred microliters of 1% orthophosphoric acid and 500 μL of precipitation mixture (1% thiobarbituric acid in 1% acetic acid) were added to 100 μL of homogenate. The resulting reaction mixture was homogenized and incubated for 15 minutes in a boiling water bath. After cooling in an ice bath, the mixture was centrifuged at 3500 rpm for 10 min. The absorbance of the supernatants was read at 532 nm against the blank. Lipid peroxidation was calculated on the basis of the MDA molar extinction coefficient and expressed in micromoles of MDA per gram of tissue using the Beer-Lambert law.

#### Determination of catalase

The catalase activity was assessed in plasma and tissues by the method of [30], with some modifications. Twenty-five microliters of plasma homogenate or tissues were added to 375 μl of phosphate buffer pH 7.4. Then 100 μl of H_2_0_2_ (50 mM) was introduced into the test tubes. One minute later, 1 ml of potassium dichromate (5%) prepared in 1% acetic acid was introduced into the reaction medium. The tubes were then incubated for 10 minutes in a boiling water bath and then cooled in an ice bath. The absorbance was recorded at 570 nm using the Schimadzu 1501 spectrophotometer, Japan. The enzymatic activity of catalase was deduced according to the Beer-Lambert law by [31], in mmol/min per milliliter of plasma or gram of tissue.

#### Determination of peroxidase

Peroxidase activity was determined using the method described by [32]. To do this, 0.5 ml of homogenate or plasma was added to 1 ml of a solution of potassium iodide (10 mM) and 1 ml of sodium acetate (40 mM) was added. The absorbance of the sample was read at 353 nm, which indicated the amount of peroxidase. Then 20 μl of H_2_O_2_ (15 mM) was added, and the change in absorbance in 5 min was recorded. The enzymatic activity of peroxidase was expressed in μmole/min per milliliter of blood or gram of tissue according to Beer-Lambert’s law by [28].

## STATISTICAL ANALYSES

The results were expressed as mean ± Standard deviation. Statistical analysis of data was performed by one way analysis of variance (ANOVA), followed by Waller-Duncan test. P values < 0.05 were considered as significant. IC_50_ were determined using Graph pad prism software (5.0).

## RESULTS

### Effects of *Dracaena deisteliana* leaves extracts on DPPH• radical

The results of the DPPH antiradical activity of the different extracts are shown in figure 1. This figure shown that these extracts possess antiradical activities. In addition, these activities are concentration dependent for each extract tested. However, 65% hydroethanolic extract showed a higher activity than the other extracts in 200 μl/mg concentration. In addition, 55% hydroethanolic extract showed a higher activity than the other extracts in 100 μl/mg concentration; the infused showed lower antiradical activity than all extracts at 12.5 and 200 ug/ml concentrations. In general, the activity of L-ascorbic acid was significantly higher (p < 0.05) than all extracts at all concentrations.

**Figure 1:**
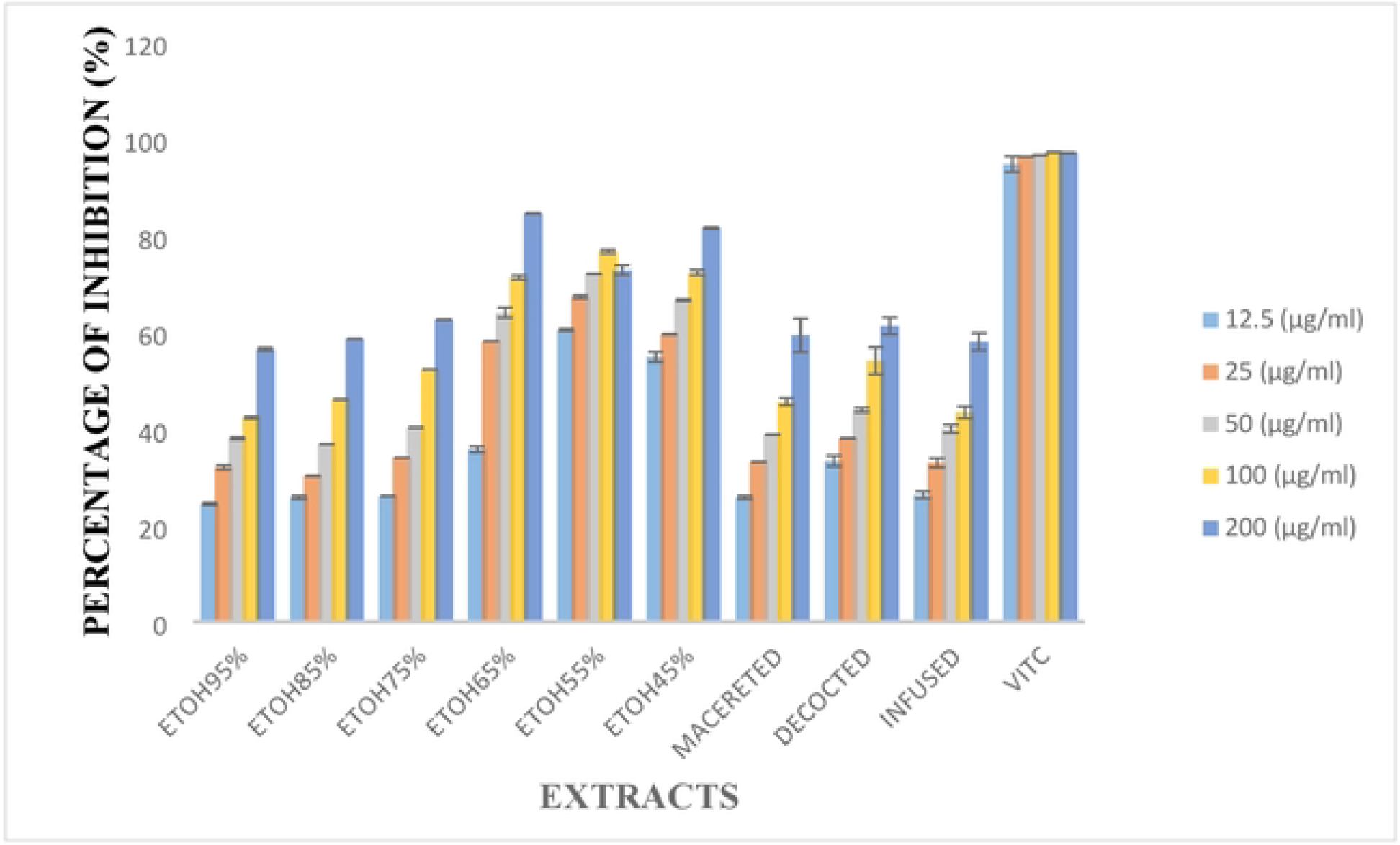
Percentage change in DPPH reduction of hydroethanolic and aqueous extracts in function of the concentration of *Dracaena deisteliana* leaves extracts. ETOH: Ethanol

### Determination of IC_50_ extracts of *Dracaena deisteliana* leaves

In order to better express the antioxidant capacity of the different extracts, inhibition percentages were used for the determination of IC_50_ (concentration needed to reduce 50% of the DPPH• radical). The IC_50_ values obtained for all extracts tested are shown in Figure 2. The 55% hydroethanolic extract had the lowest IC_50_ 11.99 μg/ml among the hydroethanolic extracts, while the decoctate had the smallest IC_50_ 96 μg/ml among aqueous extracts. The vitamin C was the most active with IC_50_ 0.01705 μg/ml.

**Figure 2:**
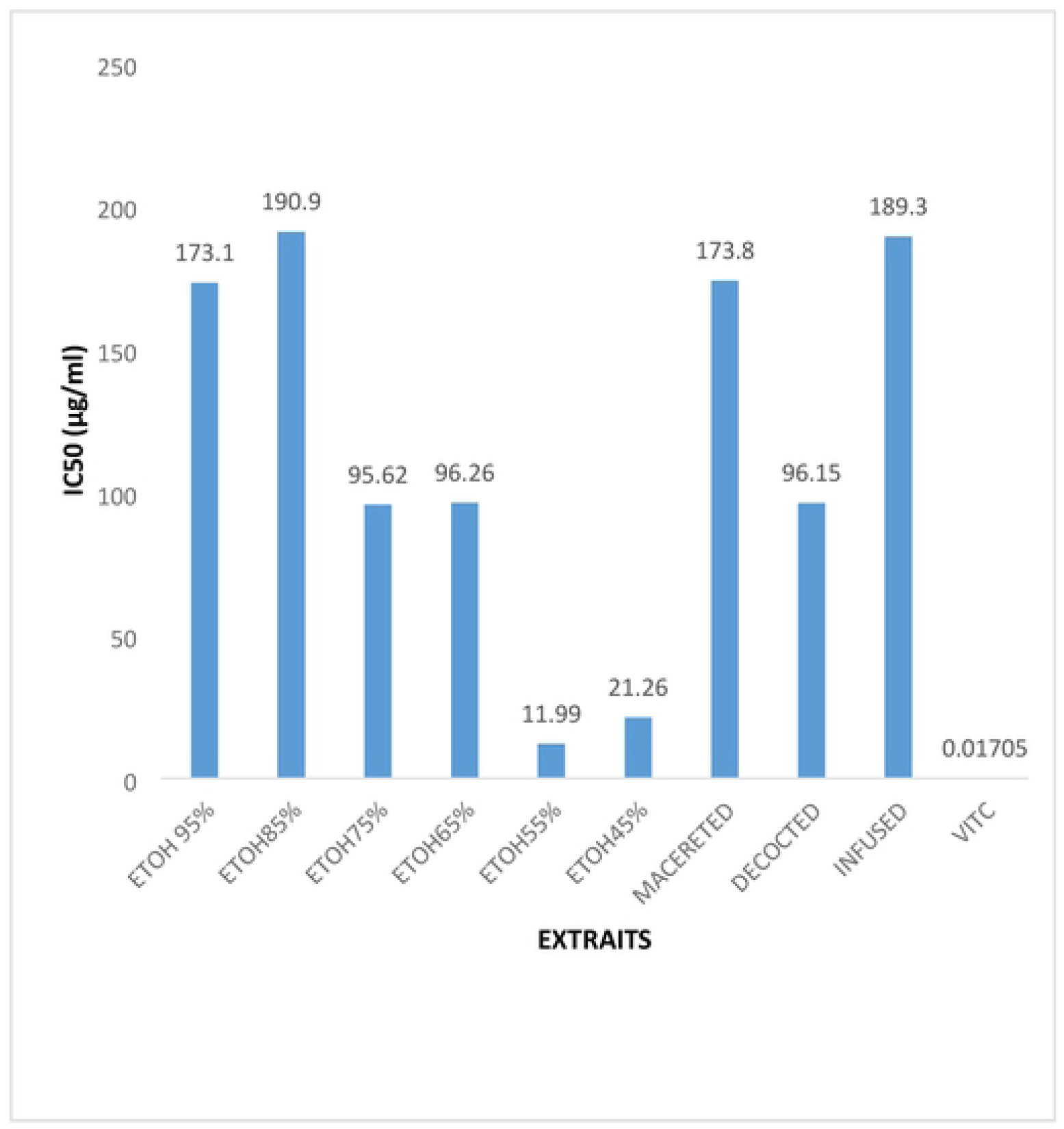
Determination of the IC_50_ of hydroethanolic and aqueous extracts in function of the concentrations of *Dracaena deisteliana* leaves. ETOH: Ethanol

### Iron reducing power (FRAP) of extracts from *Dracaena deisteliana* leaves

The results of the iron reducing power of *Dracaena deisteliana* extracts leaves are presented in Figure 3. The hydroethanolic extract 55% of the leaves of *Dracaena deisteliana* had the highest reducing power (p>0.05) compared to all extracts at concentrations of 100 and 200 μg/ml. The infused had the lowest reducing (p<0.05) power compared to all extracts at all concentrations.

**Figure 3:**
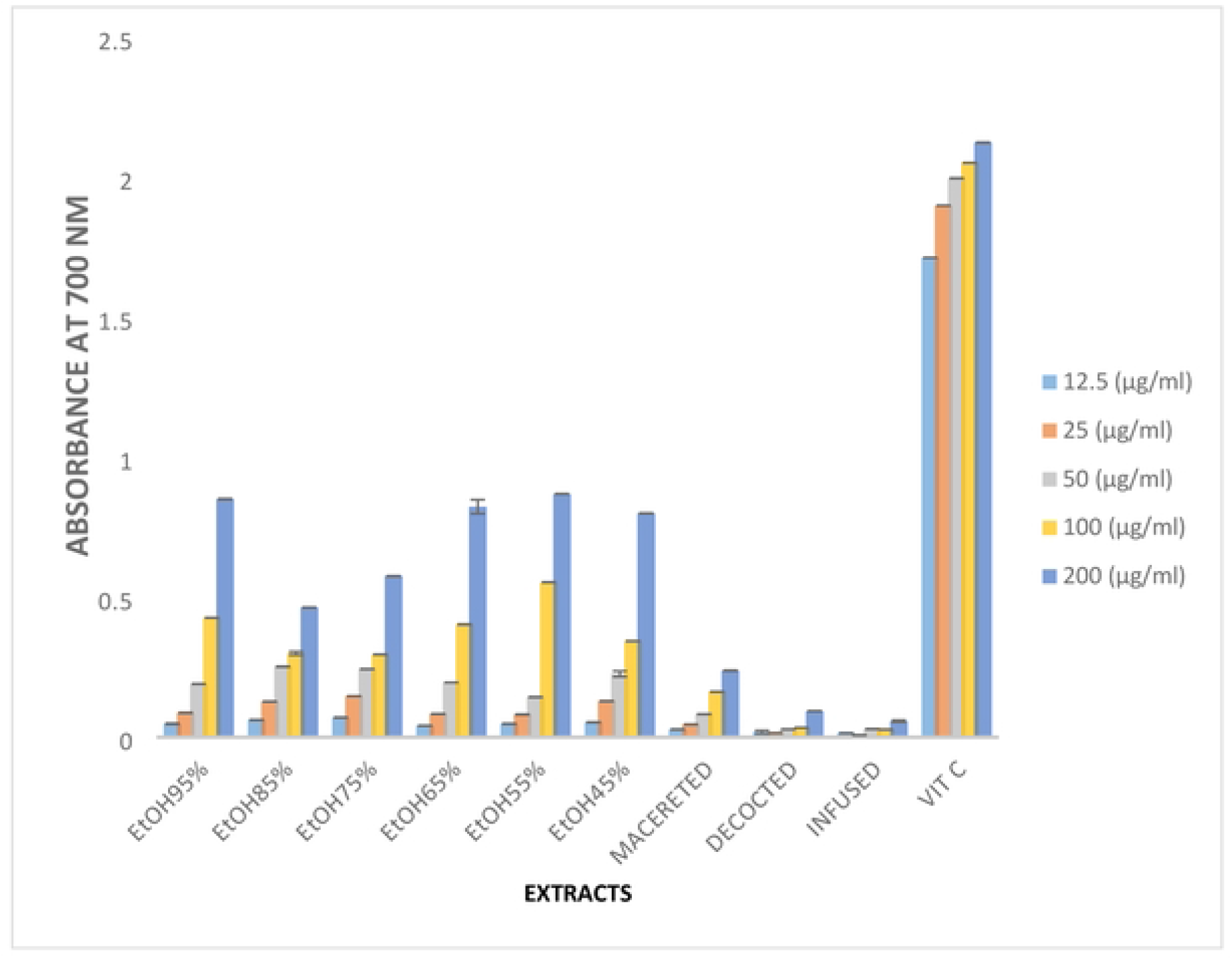
Iron reducing power of hydroethanolic and aqueous extracts of *Dracaena deisteliana* leaves. ETOH: Ethanol

### Total flavonoid content

Figure 4 shows the total flavonoid content of the extracts of the leaves of *Dracaena deisteliana*. It should be noted that the flavonoid content of the infused was significantly higher (p>0.05) than that of all other extracts. The 85% hydroethanolic extract had the lowest (p < 0.05) content compared to other extracts.

**Figure 4:**
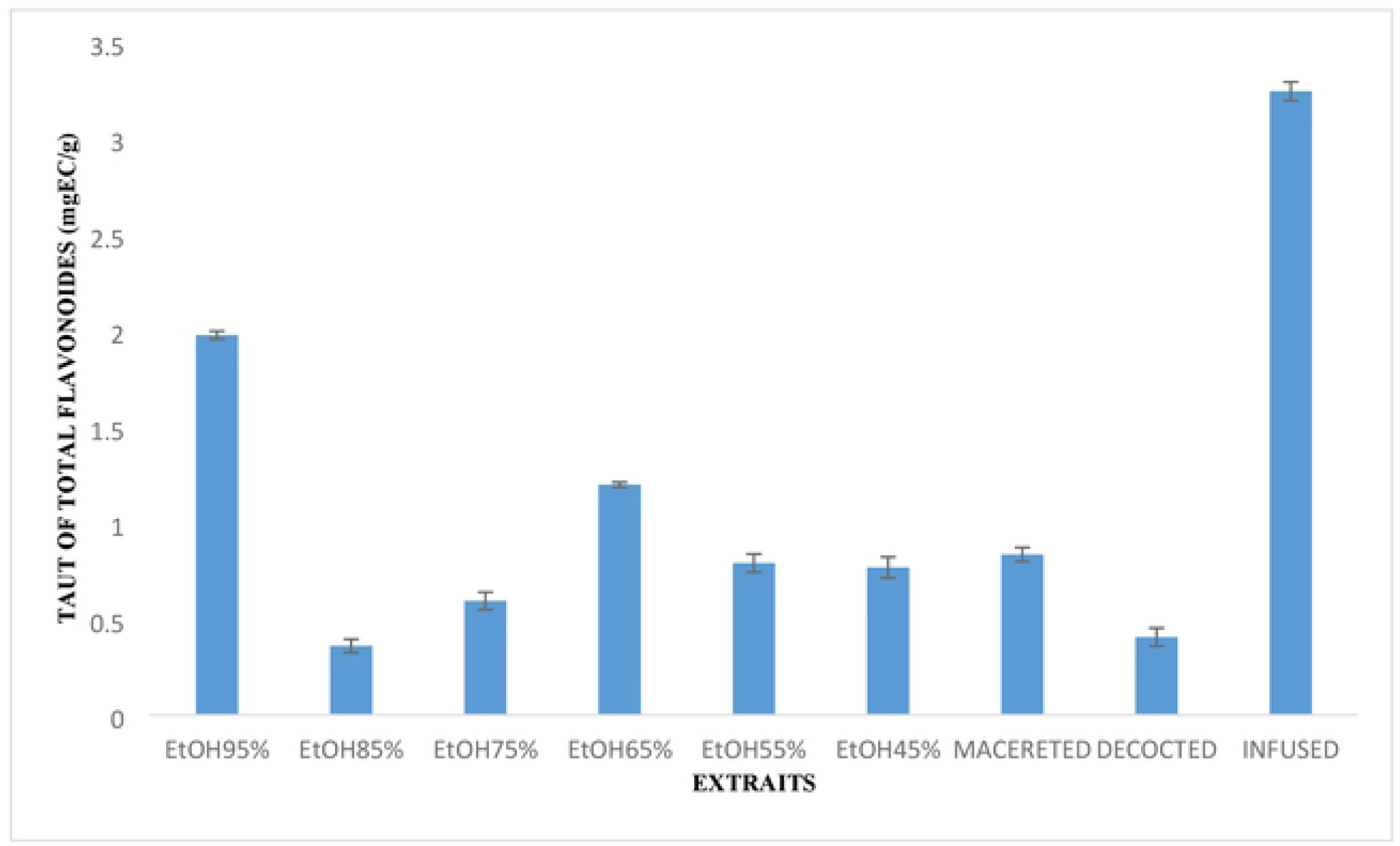
Total flavonoid content of hydroethanolic and aqueous extracts of *Dracaena deisteliana* leaves. ETOH: Ethanol

### Total phenol content

The total phenol content of the different extracts of the *Dracaena deisteliana* leaves has been determined and the results are presented in Figure 5. It should be noted that the total phenol content of the infused was significantly higher (p>0.05) than that of all other extracts, the 75% hydroethanolic extract had the lowest (p<0.05) content compared to other extracts.

**Figure 5:**
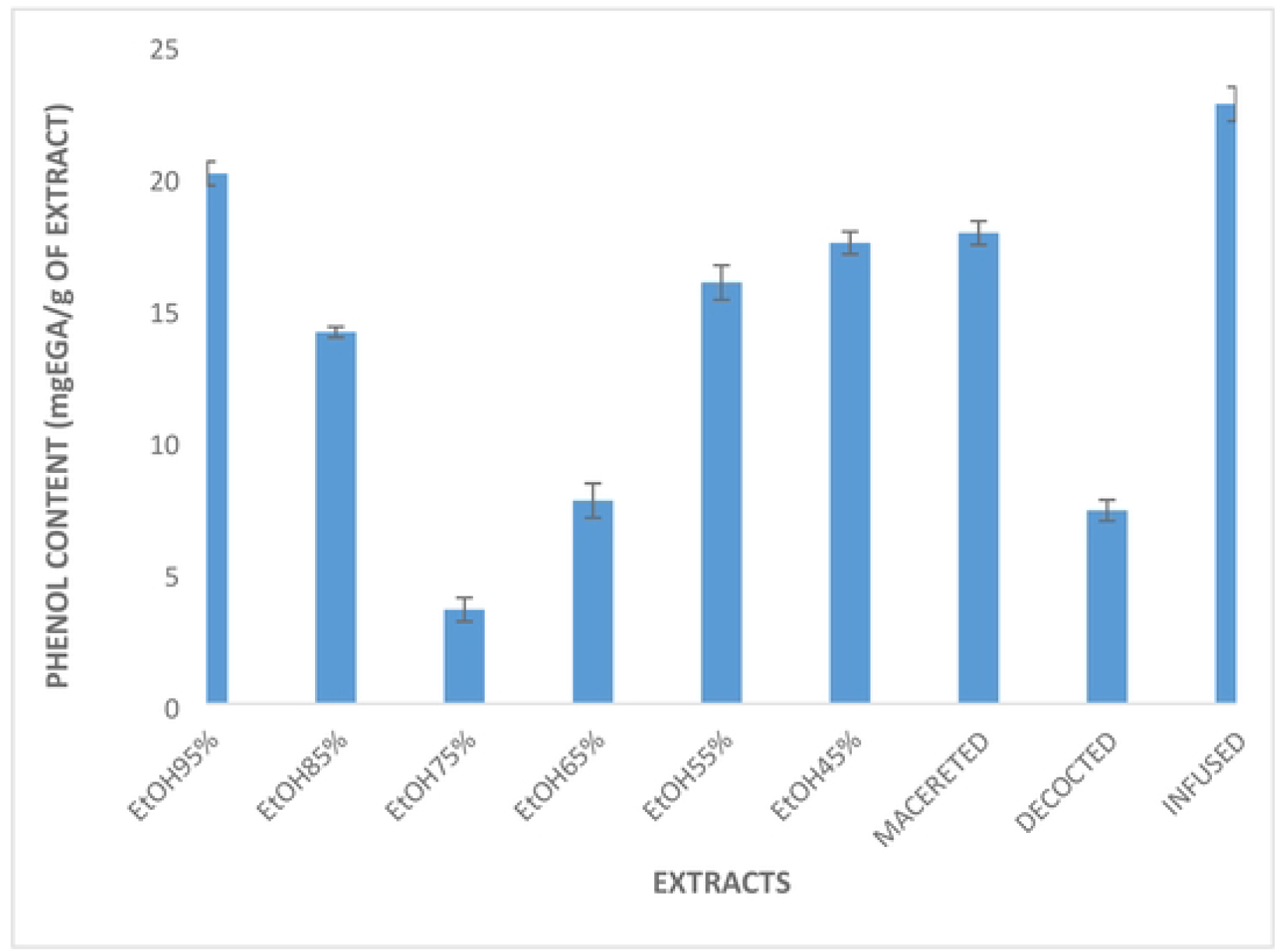
Total phenol content of hydroethanolic and aqueous extracts of *Dracaena deisteliana* leaves. ETOH: Ethanol

### *In vivo* antioxidant effect of *Dracaena deisteliana* leaves extracts

#### Effects of treatment on nitric oxide (NO) concentration

Table 1 shows the effect of treatment on nitric oxide (NO) content in the tissues of male and female rats. It shows that the infection significantly increased (p>0.05) the level of NO in all tissues of animals of both sexes compared to neutral controls. Treatment with different doses of extract resulted in a significant decrease (p<0.05) in NO levels in all organs on which the tests were performed in dose-dependent ways. Overall, treatment tends to normalize NO levels in spleen of both sexes compared to neutral (uninfected/untreated) controls in all doses.

#### Effects of treatment on malondialdehyde concentration

Table 2 shows the variation in malondialdehyde (MDA) levels in animal tissue homogenates. It shows that the infection significantly increased (p>0.05) the level of MDA in all tissues of animals of both sexes compared to neutral controls. Treatment with different doses of extract resulted in a significant decrease (p< 0.05) in the level of MDA in all organs on which the tests were performed in dose-dependent ways. Overall, treatment tends to normalize MDA levels in rats of both sexes compared to neutral (uninfected/untreated) control in all doses.

#### Effects of treatment on catalase activity

Table 3 presents the effect of treatment on catalase activity in the tissues and plasma of male and female rats. It shows that the infection induced a significant decrease (p>0.05) in the catalase activity in all tissues of animals of both sexes compared to neutral controls. Treatment with different doses of extract resulted in a significant increase (p> 0.05) the level of catalase in all organs on which the tests were performed in dose-dependent ways. Infection induced a significant decrease (p< 0.05) in the plasma level of infected and untreated male rats. However, in females this infection did not induce any significant changes in the plasma of infected and untreated rats compared to rats that were neither infected nor treated. The treatment induced a significant increase (p> 0.05) in the level of catalase in the plasma of males infected rats and treated with different doses of 55% hydroethanolic extract dosedependent.

#### Effects of treatment on peroxidase activity

Table 4 presents the effect of treatment on peroxidase activity in the tissues of male and female rats. It appears that the infection induced a significant decrease (p< 0.05) in the level of peroxidase in all tissues of animals of both sexes compared to neutral control. Overall, treatment tends to normalize peroxidase levels in rats of both sexes compared to neutral (uninfected/untreated) control in all doses.

#### Phytochemistry of *Dracaena deisteliana* extracts

Qualitative phytochemical screening of extracts from the *D. deisteliana* leaves has led to the identification of several classes of secondary metabolites. The results are grouped in Table 5. It appears from this table that flavonoid phenols, saponins and anthocyanins are present in all the extracts tested. However, triterpenoids, tannins and anthraquinones are present in all tested extracts free in the 95% extract. Sterols are only present in hydroethanolique extracts 85 and 65%. None of these extracts have alkaloids.

## Discussion

### *In vitro* Antioxidant activity

Several methods exist in the literature to assess the *in vitro* antioxidant potential of a substance. Given the complexity and diversity of oxidation processes, coupled with the structural diversity of secondary metabolites, it is recommended to use more than one method to assess the *in vitro* antioxidant activity of a given substance [33]. This explains the use of three different methods for the evaluation of *in vitro* antioxidant activity of *Dracaena deisteliana* extracts in this work. 65% hydroethanolic extract showed a higher activity than the other extracts in 200 μl/mg concentration. In addition, 55% hydroethanolic extract showed a higher activity than the other extracts in 100 μl/mg concentration. However, the infused showed lower antiradical activity than all extracts at 12.5 and 200 ug/ml concentrations. These results showed that the percentages of inhibition change according to the extracts and to their concentrations. These percentages of inhibition indicate that the extracts possess an anti-radical power. The anti-radical activity of these extracts is explained by the presence of different secondary metabolites such as flavonoids [31]; contained in extracts leaves of *Dracaena deisteliana*. Indeed, flavonoids have been shown to discolor DPPH due to their ability to produce hydrogen, and protective effects in biological systems are related to their ability to transfer protons to free radicals [34]. This is in line with the work carried out by [35] and [14]. Who showed that the antioxidant activity of *Satureja Montana* and *Terminalia avicennioides* extracts respectively is related to their richness in phenolic compounds. The anti-radical activity (DPPH.) of *Dracaena deisteliana* extracts was also expressed in IC_50_. According to [36], the antioxidant potential of a plant is divided into three groups: high when IC_50_< 20 μg/ml, moderate when 20 μg/ml ≤ IC_50_ ≤ 75 μg/ml and low when IC_50_> 75 μg/ml. Based on this theory, the IC_50_ values of *Dracaena deisteliana* extracts show that they have a high antioxidant potential. Because the 55% hydroethanolic leaves of *Dracaena deisteliana* had the lowest IC_50_ (11.99 μg/ml) among the hydroethanolic extracts while the decocted had the lowest IC_50_ (96.15 μg/ml) among the aqueous extracts. IC_50_ values showed that the 55% hydroethanolic extract of *Dracaena deisteliana* leaves has a high antioxidant potential because its IC_50_, which is 11.99 μg/ml is less than 20 μg/ml. All other extracts have an IC_50_> 75 μg/ml and therefore have a low antioxidant potential. The strong anti-radical activity of the different extracts of *Dracaena deisteliana* leaves could be explained by the strong presence of polyphenolic compounds (total phenols, anthraquinones and flavonoids) which have been highlighted via phytochemical screening. The antioxidant potential of extracts leaves of *Dracaena deisteliana* is thought to be related to the presence of phenol and total flavonoids that have been quantitatively detected in all *Dracaena deisteliana* extracts. Indeed, total phenols and flavonoids are powerful antioxidants [37]. These results suggest that the extracts tested contain free radical scavenging agents that act as primary antioxidants. For the ability of the extracts to reduce iron, the hydroethanolic extract 55% of the leaves of *Dracaena deisteliana* had the highest reducing power compared to all extracts at concentrations of 100 and 200 μg/ml. The infused had the lowest reducing power compared to all extracts at all concentrations. The activity of these different extracts could be due to the presence in high quantities of phenols and particularly flavonoids whose antioxidant potential is recognized [38]. These results are consistent with those of several authors, who have reported a positive correlation between phenolic compounds and antioxidant activity [22]; [25]; [13]). In this case, these extracts would reduce iron, preventing the Fenton reaction, and the formation of the hydroxyl radical. This hypothesis corroborates the results obtained by [39] and [11], on the effectiveness of the leaves and bark of *Drymania diandra* to stabilize the OH. radical. It is believed that this high antioxidant power is due to the high presence of phenolic compounds in *Dracaena deisteliana* extracts. A compound’s reducing capacity can be used as an indicator of its potential antioxidant activity [40]. The presence of reducing compounds leads to a reduction of Fe^3+^(ferricyanide) / to ferrous ion (Fe^2+^) [41]. Numerous studies have shown that there is a direct correlation between antioxidant activities and the reducing power of hydroéthanolic extract of *Terminalia avicennioides* [14]. The reducing properties are generally associated with the presence of reducers, whose antioxidant action has been demonstrated by reducing chain reactions by gaining hydrogen atom by the oxidizing agent. It should also be noted that reducers react with certain peroxide precursors to prevent the formation of peroxides [42, 43]. According to the phytochemical results obtained, the extracts of the leaves of *Dracaena deisteliana* contain several compounds, namely phenols, flavonoids, sterols, terpenes, tannins, saponins, anthocyanins, anthraquinones. *Dracaena deisteliana* extracts have antisalmonellal and antioxidant activities. These compounds are responsible for the ativity of Dracaena deisteliana extracts.

### *In vivo* Antioxidant activities

Antioxidant systems are normally set up in living aerobic organisms to counteract the effect of oxidative stress [44]. As part of this work, two enzymatic antioxidants (catalase and peroxidase) and two markers of oxidative stress (MDA and NO) were evaluated in the organs and plasma of rats infected with *Salmonella* Typhi ATCC 6539 and treated with the 55% hydroethanolic extract leaves of *Dracaena deisteliana*. Catalase is an enzyme that catalyzes the dismutation reaction of hydrogen peroxide, it functions as a destructive of the latter. Decreased catalase and peroxidase levels in the negative control indicates that there was an oxidative environment that led to oxidative stress in the negative control, which could result in a high bacterial load. The increase in the levels of these enzymes in positive control animals and treated groups with the different doses of extracts, shows that these extracts have the ability to suppress the oxidizing environment induced by stress in the situation of infection. This may be due to the reduction in bacterial load in the treated groups compared to the negative control [14]. This suggests that the 55% hydroethanolic extract of *Dracaena deisteliana* leaves contains antioxidant substances that neutralize reactive oxygen species, thus preventing the inhibition or destruction of catalase and peroxidase, thus promoting the protection of these organs from tissue damage induced by these reactive compounds. The results are consistent with the [45] report, which showed a significant increase in serum catalase levels in mice infected with *Plasmodium berghei* and treated with methanol extract from the stem bark of *Anogeissus leiocarpus* [46]. The current results are also similar to those of [11, 14] who showed an increase in plasma and tissue levels of catalase in rats infected with *S. typhi* ATCC 6539 and treated with 70% hydroethanolic extract of the leaves of *Adenia lobata* and *Terminalia avicennioides* respectively.

Lipid degradation leads to the formation of malondialdehyde (MDA), which is used as a marker of free radical damage [47]. The amplification of damage caused by salmonellosis or stress is related to the increase in the infiltration of neutrophils into tissues. Tissue and plasma MDA levels were significantly higher in the negative control group than in other animals. This shows that there was an oxidative environment and stress in the negative control animals, which explains the high bacterial load. This increase in infection-induced MDA level suggests increased membrane peroxidation leading to tissue damage and failure of antioxidant defense mechanism to prevent excessive free radical formation. This increase could therefore indicate a state of oxidative stress of tissue cells [29]. The significant reduction in plasma and tissue levels of MDA in the positive control and in the animals treated with the extract suggests that there was no oxidizing environment and no stress in the treated groups. This significant decrease in the level of MDA induced by the treatment in almost all organs was due to the antioxidant activity of the extract, which would either prevent the destruction of the membrane bilayer of the cells or neutralize free radicals such as hydroxyl radical (OH•) and hydrogen peroxides (H202) and known for their ability to produce tissue peroxidation of cells by attacking polyunsaturated fatty acids [48]. This may also be due to the reduction in bacterial load in treated animals compared to the negative control. These results are similar to those of [25, 11, 14] who showed that extracts of *Albizia gummifera, Curcuma longa* and *Terminalia avicennioides* respectively have protective effects on membrane lipids. The significant decrease in nitric oxide levels in the tissues of infected rats treated with the 55% hydroethanolic extract leaves of *Dracaena deisteliana* compared to the negative control indicates that the extract regulates the production of nitric oxide according to its dose compared to uninfected and untreated animals. Due to its ability to regulate nitric oxide production, this extract has an activity that can give it the potential to control salmonellosis as well as the oxidative stress that can result from it. They also chelate metal ions such as iron and copper, inhibit enzymes that produce free radicals, act as destroyers of the lipid peroxidation chain, and regulate antioxidant enzymes [49].

## Conclusion

The results demonstrate that all aqueous and hydroethanolic extracts of *Dracaena deisteliana* leaves possess *in vitro* antioxidant activities. The 55% hydroethanolic extract leaves of *Dracaena deisteliana* has both *in vitro* and *in vivo* antioxidant activity and therefore it could be used to counteract the oxidative stress generated during typhic salmonelloses. However, studies must be conducted on the toxicity of this extract in order to establish its safety.

## Supporting information

**Table 1:** Effects of different doses of treatment on the activity of nitric oxide (NO)

**Table 2:** Effects of different doses of treatment on malondialdehyde concentration as a function of extract doses

**Table 3:** Effects of different doses of treatment on catalase activity as a function of doses.

**Table 4:** Effects of different doses of treatment on peroxidase activity as a function of dose.

**Table 5:** Phytochemical composition of extracts from the leaves of *D. deisteliana*

## Acknowledgements

The authors acknowledge the Laboratory of Bacteriology of Centre Pasteur du Cameroun for providing Salmonella isolates and the Cameroon National Herbarium (Yaoundé) for plant identification. The authors also acknowledge the head of Research Unit of Microbiology and Antimicrobial Substances (Pr Kuiaté Jules-Roger).

## Author Contributions

**Conceptualization:** Hguette Bocanestine Laure Feudjio, Marie Stéphanie Chekem Goka

**Data curation:** Hguette Bocanestine Laure Feudjio, Marie Stéphanie Chekem Goka

**Formal analysis:** Hguette Bocanestine Laure Feudjio

**Funding acquisition:** Josias Djeguemtar

**Investigation:** Hguette Bocanestine Laure Feudjio

**Methodology:** Hguette Bocanestine Laure Feudjio

**Project administration:** Gabriel Tchuente Kamsu, Donatien Gatsing

**Resources:** Hguette Bocanestine Laure Feudjio

**Software:** Hguette Bocanestine Laure Feudjio

**Visualization:** Jean Baptiste Sokoudjou, Norbert Kodjio, Louis-Claire Ndel Famen,

**Writing – original draft:** Hguette Bocanestine Laure Feudjio

**Writing – review & editing:** Hguette Bocanestine Laure Feudjio

## Data Availability

No additional data are available

## Availability of data and materials

All data generated or analysed during this study are included in this published article.

## Declarations

### Ethics approval and consent to participate

This research were performed after approval by the University of Dschang-Cameroon Ethics Committee (Project N° BCH1202/FS/UDs/2018).

### Consent for publication

All authors read and approved the final manuscript

## References

1) Farr S.B., Kogoma T. (1991). Oxidative Stress Responses in *Escherichia coli* and Salmonella typhimurium. American Society for Microbiology, 55(4): 561–585.

2) Madigan M., Martinko J. (2007). Brock Biologie des Microorganismes. (11e édition) Pearson Education: Paris, France, 1046 p.

3) Ferric C.F., Vazquez-Torres A. (2002). Nitric oxide production by human macrophages: there’s NO doubt about it. American Journal of Physiology. Lung Cellular and Molecular Physiology, 282: 941–943.

4) Victor M.V., Milagros R., Monica D.L.F. (2004). Immune cells: free radicals and antioxidants in sepsis. International Immunopharmacology, 4: 327–347.

5) (5’)Rastaldo R, Pagliaro P, Cappello S, Mancardi D, Westerhof N, Losano G. Nitric oxid and cardiac function. Life Sci. 2007;81:779–93.

6) (6’)Hyun DH, Hernandez JO, Mattson MP, de Cabo R. The plasma membrane redox system in aging. Aging Res Rev. 2006;5: 209–20. 7.

7) (7’)Onanuga IO, Ibrahim RB, Amin A, Akinola OB, Omotoso GO, Ogundele MO, Tagoe CNB. Evaluation of the effects of ascorbic acid on azathioprine-induced alteration in the testes of adult Wistar rats. Int J Biol Chem Sci. 2014;8(2)-426–33.

8) Yang L, Mih N, Anand A, Park JH, Tan J, Yurkovich JT, MonK JM et al. Cellular responses to reactive oxygen species are predicted from molecular mechanisms. Proceedings of the National Academy of Sciences. 2019;116(28):14368–14373.

9) Favier A. Le stress oxydant. Intérêt conceptuel et expérimental dans la compréhension des mécanismes des maladies et potentiel thérapeutique. L’Actualité Chimique 2003; 270: 108–115.

10) Song F.L., Gan R.Y., Zhang Y., Xiao Q., Kuang L., Li H.B. (2010). Total phenolic contents and antioxidant capacities of selected chinese medicinal plants. International Journal of Molecular Sciences, 11: 2362–2372.

11) Fowa AB, Fodouop CSP, Fokunang NC, Djoueudom GF, Famen NLC, Ongbayokolak NS, Gatsing D. (2019). Antityphoid and antioxidant activities of hydroethanolic leaf extract of Adenia lobata Jacq. (Passifloraceae) on Salmonella typhi infected wistar rats. J Med Plants Stud.; 7(1):13–22

12) Kamsu G. T., Simo Tagne R., Fodouop S. P. C., Famen N.L.C., kodjio N., ekom E.S., Gatsing D. (2019). “In vitro antisalmonellal and antioxidant activities of leaves extracts of *Tectona grandis* L. F. (Verbenaceae),” European Journal of Medicinal Plants, vol. 29, no. 4, pp. 1—13,

13) Sokoudjou Jean Baptiste, Njateng Guy Sedar Singor, Siméon Pierre Chegaing Fodouop, Norbert Kodjio, Serge Secco atsafack, Alain Bertrand Fower, Merline Namékon Djiméli Gonatien Gating. (2018). In vitro antisalmonellal and antioxydant activities of Canarium schhweinfurthii stem bark extracts. Academia Journal of Medecinal Plants 6(10): 331–341 DOI; 10.15413/ajmp.2018.0166

14) Famen LCN, Talom BT, Tagne RS, Fodouop SPC, Kamsu GT, Kodjio N, Fowa AB, Gatsing D. (2021). In vivo Antioxidant Activity of Hydroethanol Extracts of Terminalia avicennioides (Combretaceae) in Salmonella typhi-Infected Wistar rat’s model. Trop J Nat Prod Res., 5(7):1185–1191. doi.org/10.26538/tjnpr/v5i7.3

15) Elodie Yamako Konack, Jean Baptiste Sokoudjou, Norbert Kodjio, Gabriel Tchuente Kamsu, Huguette Bocanestine Laure Feudjio, Jean Raphaël Kana, Donatien Gatsing. (2021). Salmonella typhimurium infection: antioxidant status of infected poultry and efficacy of treatment with Khaya grandifoliola extract. Journal of Pharmaceutical and Scientific Innovation. 10(3):72–79.

16) Aquaron, M., 2005: Les causeries en Montagne, Sabenca dela Valéia, Barcelonnette. Conférence du 18/08/05

17) Qidwai, W and Ashfaq T., 2013: Role of Garlic Usage in Cardiovascular Disease Prevention: An Evidence Based.

18) Mensor LL, Menezes FS, Leitao GG, Reis ASO, Dos Santos TC, Coube CS, Leitao SG (2001). Screening of Brazilian plant extracts for antioxidant activity by the used DPPH free radical method. Phytother. Res. 15(2): 127–130

19) Telefo, PB; L.L Lienou;M.D Ymele, M.C. Lemfack, C; Mouokeu, C.S. Goka, S.R; Tagne, F.P. Moundipa. (2011). Ethnopharmacological survey of plants used for the treatement of female infertility in Baham, Cameroon. Journal of Ethnopharmacology, 136, 178–187.

20) Jiofack, T., L. Ayissi, C. Fokunang, N. Guedje and V. Kemeuze. (2009). Ethnobotany and ohytomedicine of the upper Nyon valley forest in Cameroon. African Journal of Pharmacy and Pharmacology vol.3(4).PP 144–150, April, 2009.

21) Mensor LL, Menezes FS, Leitao GG, Reis ASO, Dos Santos TC, Coube CS, Leitao SG (2001). Screening of Brazilian plant extracts for antioxidant activity by the used DPPH free radical method. Phytother. Res. 15(2): 127–130

22) Noghogne R.L., Gatsing D., Fotso., Kodjio N., Sokoudjou. B.J., Kuiate R.J. (2015). In vitro antisalmonellal and antioxidant properties of Mangifera indica L. Stem bark crude extracts and fractions. British Journal of Pharmaceutical Research, 5(1): 29–41

23) Padmaja M., Sravanthi M., Hemalatha K.P.J. (2011). Evaluation of Antioxidant Activity of Two Indian Medicinal Plants. Journal of Phytology, 3(3): 86–91.

24) Mohammed AI, Neil AK, Shahidul IM (2013). In vitro anti-oxidative activities and gc-ms analysis of various solvent extracts of cassia singueana parts. Drug Res. 70(4): 709–719.

24a) Moncada S, Palmer RMJ, Higgs EA (1991).

25) Kodjio N., Atsafack S.S., Njateng S.S.G., Sokoudjou B.J., Kuiate R.J., Gatsing D. (2016). Antioxidant effect of aqueous extract of *Curcuma longa* Rhizomes (Zingiberaceae) in the typhoid fever induced in Wistar Rats Model. Journal of Advances in Medical and Pharmaceutical, 7(3): 1–13.

26) Ramde-Tiendrebeogo A, Tibiri A, Hilou A, Lompo M, Millogo-Kone H, Nacoulma OG, Guissou IP. (2012). Antioxidative and antibacterial activities of phenolic compounds from Ficus sue Forssk. International Journal of Biological and Chemical Sciences. 6(1):328–336.

27) Tala DS, Gatsing D, Fodouop SPC, Fokunang C, Kengni F, Djimeli MN. (2015). In vivo anti-salmonella activity of aqueous extract of Euphorbia prostrata Aiton (Euphorbiaceae) and its toxicological evaluation. Asian Pac J Trop Biomed.; 5(4):310–318

28) Griess P. (1879). Bemerkungen zu der abhandlung der HH, Weselsky, Benedikt. Ueber einige azoverbindungen. Chem Ber.12:426–8

29) Fodouop C.S.P., Gatsing D., Talom T.B., Simo T.R., Tala S.D., Tchoumboué J., Kuiate J.R. (2014). Effect of Salmonella typhimurium infection on rat’s cell oxidation and in vivo antioxidant activity of Vitellaria paradoxa and Ludwigia abyssinica aqueous extract. Asian Pacific Journal of Tropical Disease, 5(1): 38–46

30) Dimo T., Tsala D.E., Dzeufiet D.P.D., Penlap B.V., Njifutie N. (2006). Effects of Alafia multiflora stape on lipid peroxidation and antioxidant enzyme status in carbon tetrachloride-treated rats. Pharmacologyonline, 2: 76–89.

31) Chakroun A., Jemmali A., Hamed K.B., Abdelli C., Druart P. (2007). Effet du nitrate d’ammonium sur le développement et l’activité des enzymes anti-oxydantes du fraisier (Fragaria x ananassa L.) micropropagé. Biotechnologie, Agronomie, Société et Environnement. 11(2)

32) Habbu P.V., Shastry R.A., Mahadevan K.M., Hanumanthachar J., Das S.K. (2008). Hepatoprotective and antioxidant effects of argyreia speciosa in rats. African Journal Traditional Complementary and Alternative Medicines, 5(2): 158–164.

33) Alam M.N., Bristi N.J., Rafiquzzaman M. (2013). Review on in vivo and in vitro methods evaluation of antioxidant activity. Saudi Pharmaceutical Journal, 21: 143–152.

34) Musa FM, Ameh JB, Ado SA, Olonitola OS. (2016). Evaluation of phytochemical and antibacterial properties of Terminalia avicennioides crude extract against selected bacteria from diarrhoeic patients. Bajopas.;9(1):131–132.

35) Ramde-Tiendrebeogo A., Tibiri A., Hilou A., Lompo M., Millogo-Kone H., Nacoulma O.G., Guissou I.P. (2012). Antioxidative and antibacterial activities of phenolic compounds from Ficus sue Forssk. International Journal of Biological and Chemical Sciences, 6(1): 328–336.

36) Ćetković GS, Čanadanović-Brunet J, Djilas SM, Tumbas VT, Markov SL, Cetković DD. (2007). Atioxidant potential, lipid peroxidation inhibition and antimicrobial activities of Satureja Montana L. subsp. Kitaibelii Extracts. International Journal of Molecular Sciences.;8(10):1013–1027

37) Souri E., Amin G., Farsam H., Barazandeh T.M. (2008). Screening of antioxidant activity and phenolic content of 24 medicinal plants. Journal of Pharmaceutical Sciences, 16(2): 83–87.

38) Promprom W., Chatan W. (2017).GC-MS analysis and antioxidant activity of Bauhinia nakhonphanomensis leaf ethanolic extract. Pharmacognosie Journal, 9(5): 663–667.

39) Cheng A., Chen X., Jin Q., Wenliang W., John S., Yaobo L. (2013). Comparison of phenolic content and antioxidant capacity of red and yellow onions. Czech Journal of Food Science, 31(5): 501–508.

40) Palash M, Tarum K, Mitali G. (2009). Free radical scanvening activity and phytochemical analysis in the leaf and stem of Drymania diandra Blume. International Journal of Integrative Biology. 2:82–83.

41) Sousa A, Ferreira ICFR, Barros L, Bento A, Pereira JA. (2008). Antioxidant potential of traditional stoned table olives ‘‘Alcaparras”: influence of the solvent and temperature extraction conditions. LWT Food Science Technology. 41:739–745

42) Mathew S, Abraham TE. (2006). In vitro antioxidant activities and scavenging effects of Cinnamomum verum leaf extract assayed by different methodologies. Journal Food Chemical Toxicology. 44:198–206.

43) Guessom K.O, Miaffo D, Kodjio N, Oumar M, Kuiate J.R, Kamanyi A. In Vitro Antioxidant Properties of Aqueous and Methanol Extracts of the Stem Bark of Anthocleista Vogelii Planch (Loganiaceae). J Pharm Chem Biol Sci 2016; 4(2):271–280.

43a) Sokoudjou JB, Fodouop SPC, Djoueudam FG, Kodjio N, Kana JR, Fowa AB, Kamsu TG, Gatsing D. (2019). Antisalmonellal and antioxidant potential of hydroethanolic extract of *Canarium schweinfurthii* Engl. (Burseraceae) in Salmonella enterica serovar Typhimurium-infected chicks. Asian Pac J Trop Biomed 9(11):474–483.

44) Yamako-Konack E, Kamsu TG, Djenguemtar J, Sokoudjou JB, Kodjio N, Feudjio HBL, Gatsing D. (2020). In vitro antioxidant and antisalmonellal activities ofhydroethanolic and aqueous stem bark extracts of Khaya grandifoliola (meliaceae). World Journal Of Pharmacy And Pharmaceutical Sciences. 9(12):1564–1583. DOI: 10.20959/wjpps202012-17664

45) Sokoudjou JB, Fodouop SPC, Djoueudam FG, Kodjio N, Kana JR, Fowa AB, Kamsu TG, Gatsing D. Antisalmonellal and antioxidant potential of hydroethanolic extract of Canarium schweinfurthii Engl. (Burseraceae) in Salmonella enterica serovar Typhimurium-infected chicks. Asian Pac J Trop Biomed 2019; 9(11):474–483

46) Louis-Claire Ndel Famen, Benjamin Tangue Talom, Richard Simo Tagne, Gabriel Tchuente Kamsu, Norbert Kodjio, Stephen Tamekou Lacmata and Donatien Gatsing. (2020). In vitro Antioxidant Activities and Effect of Hydroethanolic and Aqueous Extracts of Terminalia avicennioides (Combretaceae) on Salmonella. Microbiology Research Journal International. 30(1): 1–14. doi: 10.9734/MRJI/2020/v30i130185.

47) Olusegun Matthew Akanbi, Akhere A. Omonkhua, Christianah M. Cyril-Olutayo, Rotimi Yemi Fasimoye. (2010). The antiplasmodial activity of Anogeissus leiocarpus and its effect on oxidative stress and lipid profile in mice infected with Plasmodium bergheii. African Journal of reproductive health 14(3)209–212

48) Yadav S, Gupta VK, Gopalakrishnan A, Verma R. Antioxidant activity analysis of Ficus racemosa leaf extract. (2019). Journal of Entomology and Zoology Studies. 7(1):1443–1446.

